# First genome-wide analysis of an endangered lichen reveals isolation by distance and strong population structure

**DOI:** 10.1101/237164

**Authors:** Jessica L. Allen, Sean K. McKenzie, Robin S. Sleith, S. Elizabeth Alter

## Abstract

Lichenized fungi are evolutionarily diverse and ecologically important, but little is known about the processes driving diversification and genetic differentiation in these lineages. Though few studies have examined population genetic patterns in lichens, their geographic distributions are often assumed to be wholly shaped by ecological requirements rather than dispersal limitations. Furthermore, while their reproductive structures are observable, the lack of information about recombination mechanisms and rates can make inferences about reproductive strategies difficult. Here we investigate the population genomics of *Cetradonia linearis*, an endangered lichen narrowly endemic to the southern Appalachians of eastern North America, to test the relative contributions of environmental factors and geographic distance in shaping genetic structure, and to gain insights into the demography and reproductive biology of range restricted fungi. Analysis of genome-wide SNP data indicated strong evidence for both low rates of recombination and for strong isolation by distance, but did not support isolation by environment. Hindcast species distribution models and the spatial distribution of genetic diversity also suggested that *C. linearis* had a larger range during the last glacial maximum, especially in the southern portion of its current extent, consistent with previous findings in other southern Appalachian taxa. These results contribute to our understanding of intrinsic and extrinsic factors shaping genetic diversity and biogeographic patterns in *C. linearis*, and more broadly, in rare and endangered fungi.

## Introduction

Symbiotic fungi, including lichenized species, represent some of the most ecologically important radiations on earth. However, the processes shaping genetic differentiation and gene flow in these groups remain poorly understood. Historically, two major assumptions have shaped hypotheses about symbiotic fungal population structure and evolution. First, because most fungi produce very small spores, their distribution is thought to be limited primarily by ecological suitability rather than geographic distance (O’Malley 2007). Second, species in which no sexual reproductive structures have been observed are assumed to reproduce only asexually (Taylor et al. 2015). Phylogenetic and population genetic studies have already challenged these assumptions in fungi that do not form lichens. For instance, in the common and widespread fungus *Suillis brevipes* there is evidence for both isolation by distance (IBD) and adaptation of coastal populations to saline environmental gradients (Branco et al. 2015). Species in *Saccharomyces* show varying levels of geographic structure in their genetic differentiation, with *S. paradoxus* showing clear evidence of IBD and *S. cerevisiae* showing much less geographic structure (Liti et al. 2009). Taylor et al. (2015) reviewed the literature on clonal reproduction in fungi, concluding that numerous species showed evidence for recombination regardless of observed reproductive structures. Additional examples of similar observations, indicating that fungal reproduction and population genetics are more complex than previously expected, have been derived from population genetic and genomic data in other groups of non-lichenized fungi (see review in Grünwald et al. (2016) and Peter and Schacherer (2016)).

Lichens are a major group of fungi that form obligate symbioses with algae and/or cyanobacteria and comprise >20% of all ascomycetes (Lücking et al. 2016). Despite their ecological importance and conspicuous abundance in many terrestrial ecosystems, relatively few taxa have been studied with traditional population genetics methods, and no published studies have used a genome-wide approach to assess gene flow or other population-level attributes. To date, most population genetics studies of lichens have been conducted on *Lobaria pulmonaria* and its algal photobiont *Dictyochlorpsis reticulata* using microsatellite markers (Widmer et al. 2010, Dal Grande et al. 2010, Nadyeina et al. 2014). These studies have shown that *L. pulmonaria* frequently disperses short distances via lichenized propagules (bundles of algae and fungi), infrequently disperses long distances via sexually produced fungal spores (Werth et al. 2006), and that there is evidence of adaptation and population isolation on small spatial scales (Nadyeina et al. 2014). Population genetic patterns in another lichenized fungus, *Xanthoria parietina*, based on RAPD-PCR markers, contrast starkly with the findings for *L. pulmonaria;* in the former, high genetic diversity and very few clones were found within small areas, even among adjacent individuals (Itten and Honneger 2010). The pattern recovered in *X. parietina* is similar to a study of *Parmelina carporrhizans* based on microsatellite loci, where high rates of migration were recovered among populations, except for isolated island populations (Alors et al. 2017). A microsatellite-based study on *Parmotrema tinctorum* and its algal symbiont found that most dispersal was clonal over short distances, similar to *L. pulmonaria*, but still found evidence for high rates of sexual reproduction in the fungus (Mansournia et al. 2012). While highly detailed, these studies of lichen population genetics represent only a fraction of this diverse group of fungi that have evolved an obligate symbiotic lifestyle at least seven times independently throughout the fungal tree of life (Schoch et al. 2009), and occupy every terrestrial ecosystem from the poles to the tropics (Brodo et al. 2001). Microsatellites have recently been developed for a broader diversity of lichenized fungi (Magain et al. 2010; Devkota et al. 2014; Nadyeina et al. 2014; Lindgren et al. 2016; Lutsak et al. 2016), however these tools have not yet been utilized for population genetic analyses in lichens.

Population genomics is a promising approach to rapidly advance our knowledge of population biology in lichens as it circumvents difficulties associated with developing species-specific markers, especially since lichens are notoriously difficult and slow to culture (Crittenden *et al.* 1995). Of the domains of eukaryotic organisms, fungi are one of the most amenable to genomic studies due to their generally small, compact genomes (Gladieux et al. 2014). Population genomics studies have already added substantial depth and breadth to the knowledge of the basic fungal biology, allowing researchers to address questions that were once intractable about life history and evolution of reproductive systems. For example, fungi that have only been observed reproducing asexually show genomic evidence for sexual reproduction (Tsai et al. 2008; Stefanini et al. 2016); speciation through homoploid hybridization has been shown to occur rapidly, at least in yeast (Leducq et al. 2016); and Glomerales, arbuscular mycorrhizal fungi, have highly flexible levels of ploidy in the heterokaryotic cells within species (Wyss et al. 2016). Applying these methods to lichenized fungi holds great promise to rapidly advance knowledge of lichen population biology.

The rock gnome lichen *(Cetradonia linearis)* is one of two fungal species protected by the Endangered Species Act in the United States (USFWS 2013), and one of eight lichens on the IUCN Red-List (Allen et al. 2015). It is narrowly endemic to the Southern Appalachians of eastern North America, where it is known from ~100 populations, most of which are located in western North Carolina (USFWS 2013). It forms colonies on rocks either on exposed cliffs at high-elevations or on large boulders in mid-to high-elevation streams. *Cetradonia* is a monotypic genus, whose position as the earliest diverging member of the widespread and ecologically important Cladoniaceae makes its study essential for addressing hypotheses of evolution in this family (Wei and Ahti 2002; Zhou et al. 2006). It forms colonies of simple to branched squamules with black apothecia and/or pycnidia, reproductive structures, frequently produced at the tips (Fig. 1). Despite having been protected by the Endangered Species Act for over 20 years, little is known about *C. linearis* beyond its distribution (USFWS 2013). Currently, nothing is known about dispersal or population genetic structure in this species.

**Figure 1.**
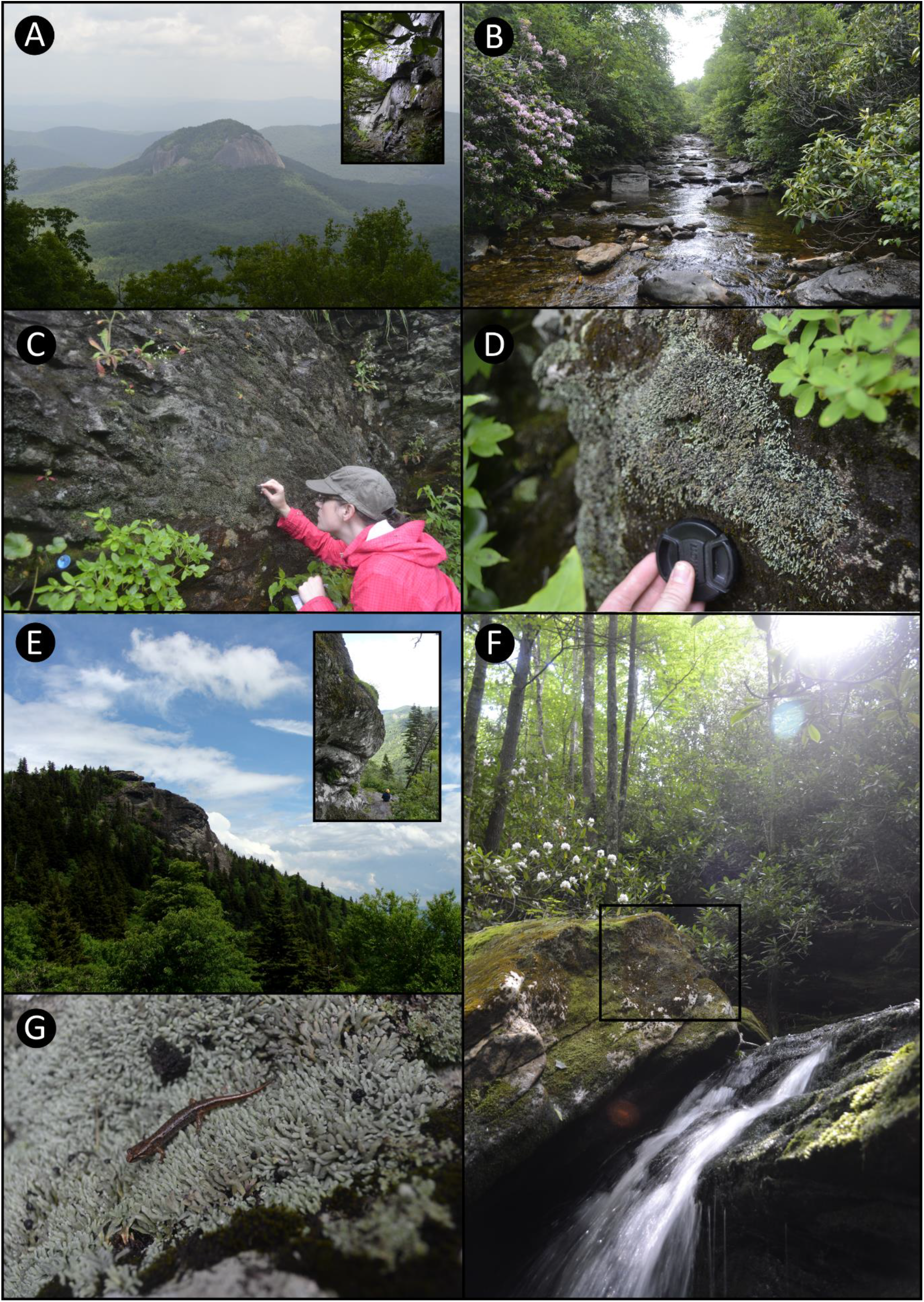
Morphology, habit, and habitat of *Cetradonia linearis*. A) Large granite dome where species occurs at base of large rock faces, inset shows seeping rock faces where the species occurs; B) Stream habitat where species occurs frequently on scattered rocks and boulders throughout; C) Large boulder face covered in the species illustrating sampling protocol using sterile forceps; D) Fertile colony on mossy boulder in stream; E) Large rock outcrop hosting colonies of the species, inset shows view from *Cetradonia linearis* perspective; F) Waterfall populations are very abundant, one colony outlined by black box; G) Colony displaying apothecium and potential zoochory event.

In this study, we tested three hypotheses concerning population-level processes in *Cetradonia linearis*: 1) most reproduction and dispersal occurs through clonal processes, 2) isolation by distance is the major force shaping the genetic differentiation, while ecological adaptation plays a minor role, and 3) the southern portion of its current extent represents a major refugium during the Pleistocene glaciation. To test these hypotheses, low-coverage, whole genome shotgun sequencing was used to generate large-quantities of genomic data from samples throughout the species’ range. The resulting genome-wide single-nucleotide polymorphisms (SNPs) were used to measure genetic diversity, recombination, and clonality. Population genetic structure, connectivity, and evidence for isolation by environment were also investigated. This study is the first assessment of population genomics in a lichen, providing a baseline for comparison in this group of organisms, along with valuable information for the continued conservation of the endangered rock gnome lichen.

## Methods

### Study System, Sampling, and Sequencing

Samples were collected from 15 sites throughout the geographic and ecological range of *Cetradonia linearis* (Fig. 2). At each site two to three squamules were taken from up to ten distinct colonies using surface sterilized forceps. Squamules were placed into 1.5 mL Eppendorf tubes, set out to air dry for 24 hours, then stored at −40°C. Samples were washed with acetone and DNA was extracted using the Qiagen DNeasy Plant Mini Kit with the cell lysis stage extended for 4-6 hours. Thirty-two samples were chosen for sequencing based on DNA quality and yield, while maintaining the geographic and ecological breadth of samples. Sequencing was conducted at the Rockefeller University Genomics Resource Center. Libraries were prepared with the Nextera XT kit and sequenced on the Illumina Next Seq platform in Mid Output on 150 bp paired end read mode. All samples were sequenced at roughly equal coverage, except one sample from the Balsam Mountains, B224, which was sequenced at 5x higher coverage for assembly of a reference genome.

**Figure g2.**
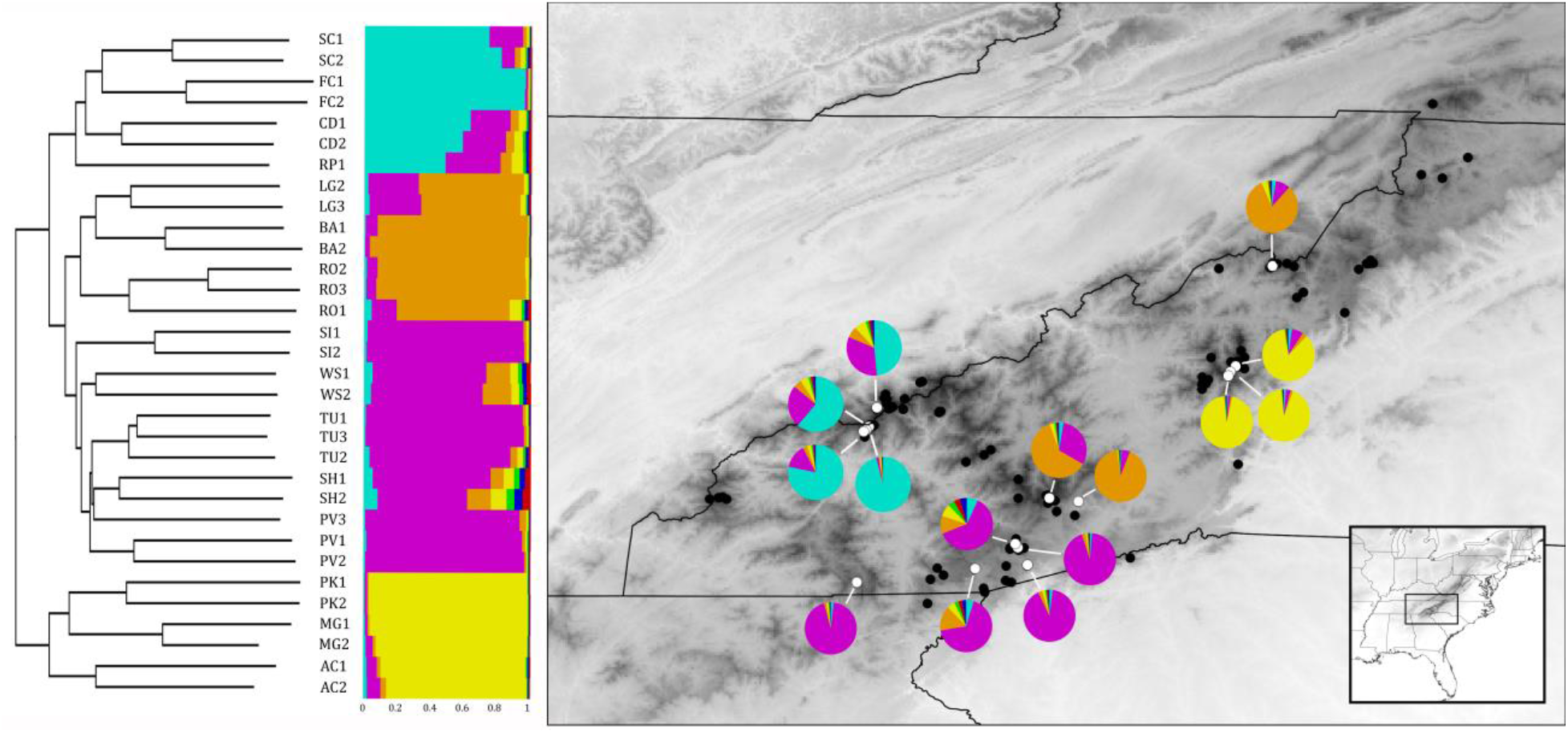
Population structure of *Cetradonia linearis.* Left: Neighbor-joining tree showing hierarchical clustering of all sampled individuals. Middle: Proportional cluster belonging of each individual sampled as inferred by InStruct. Right: Distribution of *C. linearis* and average proportional cluster belonging for each sampling site.

### Quality Filtering, Genome Assembly, and Annotation

A reference genome was assembled from sample B224 and annotated after strictly filtering contaminating reads (see below). B224 reads were trimmed, adapters were removed, and overlapping read-pairs combined using cutadapt and FLASH (Magoč and Salzberg 2011; Martin 2011). Read pools for all other samples were trimmed, adapters were removed, and overlapping read-pairs combined using FLASH and Trimmomatic v 0.36 (Bolger et al. 2014).

An initial assembly of B224 was built using Minia with a kmer size of 75 and an abundance minimum of 3 (Chikhi and Rizk 2013). To filter out contaminants (including algal symbionts) the Blobology workflow and perl scripts were used (Kumar et al. 2013). Specifically, a random subset of 15,000 contigs longer that 250 bp were subjected to homology search using megablast against the non-redundant nucleotide database from Genbank and the e-value cutoff was set to 1e-5. Based on these plots (Supplementary Fig. 1) contigs with GC content >0.6 and coverage <5 were pooled to form a set of contaminant contigs. Then, all B224 reads were aligned to the contaminant contigs using bowtie2, and all reads that did not align to the contaminants were retained for reassembly. The final assembly was built using Abyss with the paired-end read setting and a kmer size of 41 (Simpson et al. 2009). All resulting contigs shorter than 500 bp were removed from the dataset before further analyses. Genome annotation was conducted using the MAKER pipeline (Cantarel et al. 2008). SNAP was used for the ab-initio gene predictor, and protein homology evidence was drawn from *Aspergillus niger* ATCC 1015 v4.0, *Cladonia grayi* Cgr/DA2myc/ss v2.0, and *Cochliobolus heterostrophus* C5 v2.0 (Andersen et al. 2011; Ohm et al. 2012; Condon et al. 2013; McDonald et al. 2013; Leskovec and Sosič 2016). For a final filtering step, all genes were blasted against the *A. niger*, *C. grayi*, and *C. heterostrophus* gene sets. Contigs were only kept for downstream analysis if the gene with the highest-scoring blast hit matched most closely with a *C. grayi* gene. Because the *C. grayi* genome was assembled from pure culture of the fungal symbiont, this was an additional step that filtered any remaining contaminants from the genome. Finally, we conducted homology searches for both mating-type idiomorphs (MAT1-2 and MAT2-2) in the genome and all sampled read pools (Supplementary Text).

### SNP Calling and descriptive statistics

Single-nucleotide polymorphisms were called for all sequenced samples using the annotated contigs as a reference genome for the fungal component. First, we used bwa (Li and Durbin 2009) to align the reads to the contigs. Then, to call the SNPs from this alignment FreeBayes was used with the ploidy set to 2, minimum alternate fraction set to 0.9 (Garrison and Marth 2012). The ploidy was set to two because all samples were fertile, thus there were potentially two genetic individuals present (we also conducted the same analyses with the ploidy set to one, without a change in results). Average nucleotide diversity were calculated using VCFtools (Danecek et al. 2011). Linkage distance was calculated between all sites using PLINK 1.9 and plotted in R with the non-linear least squares smoothing function implemented for the trendline to create a linkage decay chart (Gaunt et al. 2007). Linkage disequilibrium was corrected for using the R package poppr with a threshold of 0.2 and a 1 Kb sliding window (Kamvar et al. 2014). Then, pairwise F_st_ was calculated among all sampling sites and populations with the linkage disequilibrium corrected dataset using BEDASSLE (Bradburd et al. 2015), a Bayesian inference program that estimates the relative influence of ecological and geographic distance on genetic distance. This program automatically excludes sites with missing data when calculating F_st_ (Bradburd et al. 2015). The dataset was checked for clones by searching for multi-locus genotypes that are >95% identical to account for sequencing and SNP calling errors using the mlg.filter function in the R package popper with a threshold of 0.05 (Kamvar et al. 2014).

### Statistical Analyses

First, the relationships among populations were explored to determine if there were phylogenetic signals for each distinct sampling site and mountain range, and at what spatial scale the relationships were clearest. Three methods were used to explore population structure: InStruct, Discriminant Analysis of Principle Components (DAPC), and a neighbor-joining tree. InStruct is a Bayesian clustering program that infers self-fertilization rates and clusters individuals into subpopulations using a Markov chain Monte Carlo (MCMC) algorithm (Gao et al. 2007). InStruct was run with K values between one and 10, with five independent chains per K value. Each chain was run with a 100,000 iteration burn-in period, followed by 10,000 iterations after burn-in. Mode 2 was used to infer subpopulation structure and selfing rate. The run with the highest DIC was chosen to represent the population structure and inferred selfing rate. We also implemented DAPC, a multivariate approach to identifying genetically distinct clusters of individuals, with the clustering algorithm implemented to define groups resulting in 10 genetic clusters chosen based on the BIC (Jombart et al. 2010). This method was specifically designed to cope with the large quantity of next generation sequencing data and implemented in R through the package adegenet 2.0 (Jombart 2008). An unrooted neighbor-joining tree was also built to infer the relationships among individuals based on bitwise distances using the R package ape (Paradis et al. 2004).

The influence of geographic and ecological distance on genetic distance was investigated using two approaches. First, a partial Mantel test with 10,000 permutations was used to test for correlation between genetic distance, measured as pairwise Fst, and geographic distance (Euclidean distance in kilometers), and a set of four environmental variables. The set of four environmental variables were habitat (boulder in stream vs. exposed rock outcrop) and three non-colinear variables from the widely-used Worldclim dataset: mean temperature of wettest quarter (BIO8), mean temperature of warmest quarter (BIO10), and annual precipitation (BIO12) (Hijmans et al. 2005). These last three variables were retained after removing all correlated climatic variables from Worldclim (see Species Distribution Modeling, below). The habitat variable was based on field observations of the species made while collecting samples. Second, a Bayesian approach as implemented in BEDASSLE was used to estimate the contributions of geographic and ecological distance to genetic distance (Bradburd et al. 2013). The same set of ecological and geographic distance variables were used as input data, along with allele sample sizes and frequencies in all samples. An initial Bayesian analysis, run for 1 million generations, indicated that the effect size of BIO8 and BIO12 were very close to zero, and these were removed from the dataset. A second analysis was run retaining the habitat and BIO10 as environmental variables for 5 million generations with a sample frequency of 10. A third analysis was conducted retaining only the habitat as the environmental variable and was run for 10 million generations with a sample frequency of 10. Trace plots were examined for convergence, and mean marginal densities and 95% confidence intervals calculated for αE:αD for each environmental variable with the first 50% of generations treated as burn-in and removed.

### Reproductive Morphology

To determine whether reproductive structures were observable in *Cetradonia* specimens, fertile samples were dissected to search microscopically for trichogynes, specialized hyphae that receive spermatia (conidia) to begin sexual reproduction, and fertile apothecia, structures that produce fungal spores that are produced through meiosis. Thin sections of five apothecia from three sampling sites were cut by hand with a razor blade through apothecia and mounted on slides. Sections were stained with phloxine and cleared with potassium hydroxide before examination under a compound microscope.

### Species Distribution Modeling

Species distribution modeling was used to investigate if the sites with the highest genetic diversity were located in an area that was likely a refugium during the Last Glacial Maximum (LGM). To model past distributions, we first built a species distribution model (SDM) to predict the probability of a species’ presence across the landscape for the present, then projected this SDM to past climates. Species distribution modeling was conducted using Maxent v. 3 (Phillips et al. 2006, Phillips and Dudik 2008) after steps were taken to reduce sampling bias and calibrate the model. First, localities were thinned by a 5 km radius to reduce sampling bias by randomly excluding one of two localities when they fell within that radius, as implemented in the R package SpThin (Aiello-Lammens et al. 2015). After thinning, 42 out of the 101 original localities were retained and used for all further analyses. The worldclim dataset of 19 bioclimatic variables was used for the environmental data at 10 arc second resolution for the present and 2.5 arc minutes for the last glacial maximum. All autocorrelated variables were first removed, leaving mean temperature of wettest quarter (BIO8), mean temperature of warmest quarter (BIO10), and annual precipitation (BIO12) (Hijmans et al. 2005). These three variables were clipped to the extent of the species known range, with a small buffer, for environmental variables from the present and LGM. Two modeling parameters were tuned to identify the best level of complexity for the model: feature classes define the allowed shape of the environmental variable response curves, and the regularization multiplier controls for complexity, with higher values increasingly penalizing complexity (Scheglovitova and Anderson 2013). The best modeling parameters were chosen based on the Akaike Information Criterion corrected for sample size (AICc) (Warren and Seifert 2011). Model tuning was implemented using the R package ENMEval with the ‘blocks’ setting (Muscarella et al. 2014). The final model was built and projected using all thinned localities with the regularization multiplier set to 3.5 and linear, quadratic, and hinge response curves allowed.

## Results

High-coverage, whole-genome shotgun sequencing of one individual of *Cetradonia linearis* was obtained and used to assemble a reference genome. Whole-genome shotgun sequencing of 31 additional individuals were mapped to this genome, and the resulting SNPs were used to infer the population structure, biogeographic history, and mating system of the species.

### Cetradonia linearis Reference Genome

Multiple steps of stringent quality and contaminant filtering resulted in the production of a high-quality, partial reference genome. The original read pool from the sample used to create the reference genome, sample B224, contained 55 million reads, for a total of 16 Gb. The mean PHRED quality score was 33. After trimming, filtering for low quality base calls, and merging paired ends there were 44 million merged reads (where two paired-end reads overlapped and merged into a single sequence) with a total of 5.6 Gb of sequence, and 8.3 million read pairs that did not overlap with a total of 2.1 Gb of sequence. The initial assembly using Minia built 32,669 contigs. When contigs under 500 bp were excluded, the total assembly length was 105.5 Mb and the N50 was 3,814 bp. After filtering contaminants, 41.9 million merged reads with 5.4 Gb of sequence remained, as well as 7.6 million paired-end read with 2.0 Gb of sequence. These filtered reads were then assembled, which built 17,199 contigs with a total length of 40.0 Mb and an N50 of 6,093. This assembly was then annotated with protein homology data from *Aspergillus niger* ATCC 1015 v4.0, *Cladonia grayi* Cgr/DA2myc/ss v2.0, and *Cochliobolus heterostrophus* C5 v2.0 and ab-initio prediction using SNAP. Then, only contigs for which the annotated gene with the best blastp score against *C. grayi* and *A.niger* proteins most closely matched *C. grayi* were retained for the final reference genome to be used in all downstream analyses. This reference genome was comprised of 2,703 contigs with a total length of 19.5 MB, a contig N50 of 10,095 bp, and an average coverage of 54.7 X. CEGMA (Parra et al. 2007) analysis of conserved gene content showed that 74% of universally conserved genes are present in our assembly, suggesting that our assembly is approximately 74% complete. Consistent with this, our assembly was 53-70% as large as the three genomes available for other species in the Cladoniaceae (28 Mb-37 Mb; Armeleo and May 2009; Park et al. 2013). The MAT1-2 idiomorph was located in the reference genome, and in 14 of 32 read pools (Supplementary text).

No MAT1-1 genes were located in any samples. Because only one mating type was discovered we preliminarily determine that the mating system of *C. linearis* may be homothallic, and specifically unisexual (Wilson et al. 2015).

### Cetradonia linearis Population Structure

To call SNPs, all read pools were aligned to the reference genome. A total of 126,662 SNPs were identified. After correcting for linkage disequilibrium 10,026 SNPs remained. This large reduction in SNPs after correcting for linkage disequilibrium suggests a low rate of recombination. Examination of the linkage decay plot further supports the hypotheses of low recombination rates, as the linkage between sites never falls below 0.2 (Fig. 3). In the dataset used for subsequent analyses the average SNP coverage was 66%, and the coverage per population ranged from 52.6-98.6% (Table 1). Nucleotide diversity (π) within sampling sites ranged from 0.084 for one site in the Great Smoky Mountains, to 0.18 for one site in the Black Mountains. When the samples were grouped by mountain range, nucleotide diversity ranged from 0.148-0.338 (Table 1). Pairwise F_st_ values between sites ranged from 0.312 to 0.730 (Supplementary Table 1).

**Figure 3.**
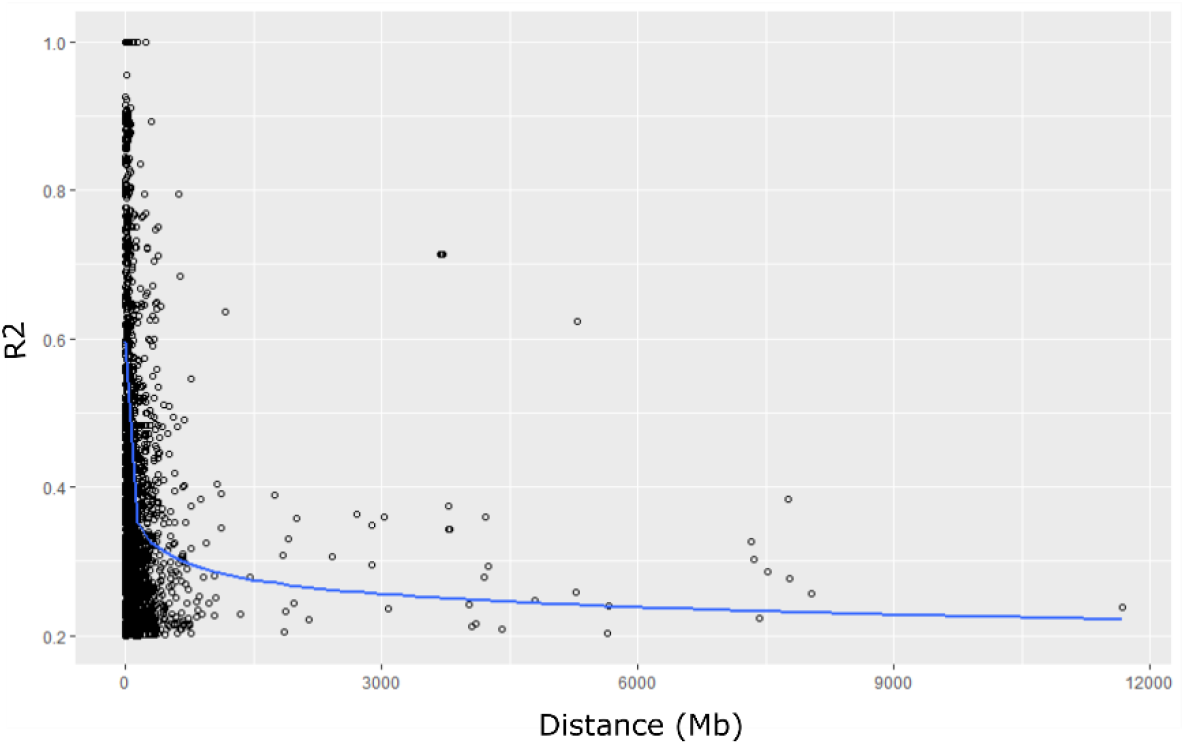
Linkage decay plot for *Cetradonia linearis.*

**Table 1.**
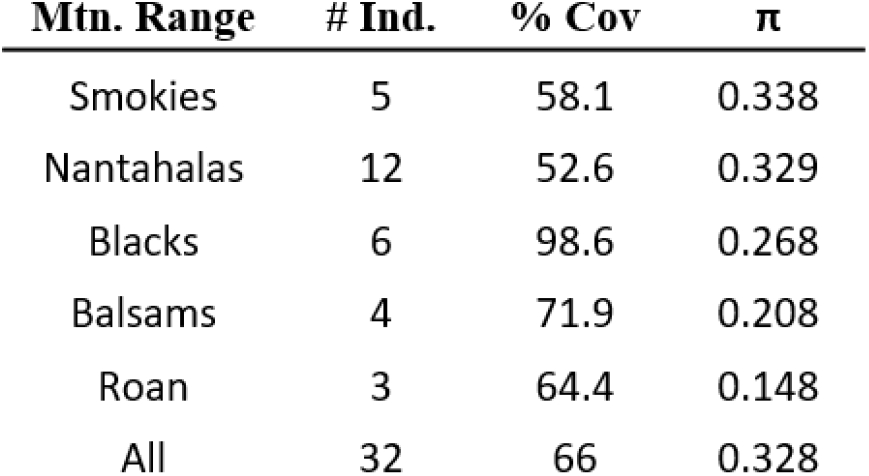
Site names, mountain range, individuals sampled, average percent SNPs covered for each site/mountain range (% Cov), and average nucleotide diversity for all sampled sites and mountain ranges.

Population structure was first explored through relational analyses. The unrooted NJ tree recovered distinct, mutually exclusive groups that corresponded to distinct mountain ranges (Fig. 2). Sampling sites also largely formed mutually exclusive groups, except PV. The one PV sample that did not cluster with other PV samples formed a group with SH, a site that was only 1.5 km downstream. The InStruct analysis chain with the highest DIC found seven clusters (Fig. 2). Four primary clusters were evident in the results: one that included all samples from the Great Smoky Mountains, one from the southern Nantahalas, one that included the Balsam Mountains and Roan Mountain, and one that included all samples from the Black Mountains (Fig. 2). The mean posterior distribution of selfing rates averaged 0.59, with the mean, followed by the variance, for each cluster inferred as follows: 1 = 0.475 (0.085), 2 = 0.511 (0.087), 3 = 0.585 (0.074), 4 = 0.606 (0.015), 5=0.620 (0.016), 6 = 0.665 (0.008), 7 = 0.666 (0.014). Ten clusters were found as the most likely grouping of the samples using DAPC. Most clusters were comprised of all individuals from single sampling sites. Group four was the only one that included samples from multiple sites, for a total of 15 individuals from nine sites that included the Great Smoky Mountains, Balsam Mountains, Nantahala Mountains, and Roan Mountain (Supplement Fig. 2). Each of the three sites sampled from the Black Mountains formed their own distinct group, despite their close proximity to each other (1-173 km apart).

We tested the influence of geographic versus environmental distance on genetic distance using two methods, and both showed geographic distance as a more significant factor correlating with population structure. First, a partial Mantel Test showed a significant correlation between genetic distance, measured as pairwise Fst, and geographic distance, measured as pairwise Euclidean distance in km, where r = 0.489, and p < 0.01 (Fig. 4). There were no correlations between genetic distance and any of the environmental distances (Supplementary Table 2). The second analysis was a Bayesian approach implemented in the program BEDASSLE (Bradburd et al. 2013). Here, the relevant value is the ratio of effect size of each environmental variable versus the effect size of the geographic distance (αE:αD). The results were similar to the partial Mantel test, and geographic distance far outweighed the effect of environmental distance. Specifically, the results of the first analysis, which included both habitat and BIO10 as environmental variables, estimated the habitat αE:αD = 0.712 (95% CI = 0.544, 0.978) and the BIO10 αE:αD = 0.026 (95% CI = 0.020, 0.036). This can be interpreted as follows: the effect of 10° C mean temperature of the warmest quarter was equal to the effect of 0.026 km of geographic distance, and the effect of occurrence in different habitats was equal to 0.75 km of geographic distance. In the second analyses, where only habitat was retained as an environmental variable, the habitat αE:αD = 0.309 (95% CI = 0.011, 1.408). Hindcasting the SDM of *Cetradonia linearis* supported the hypothesis that its refugial range was located predominantly in the southern edge of its current range during the LGM (Fig.5). The quality of the SDM was high, with an AUC of 0.919.

**Figure 4.**
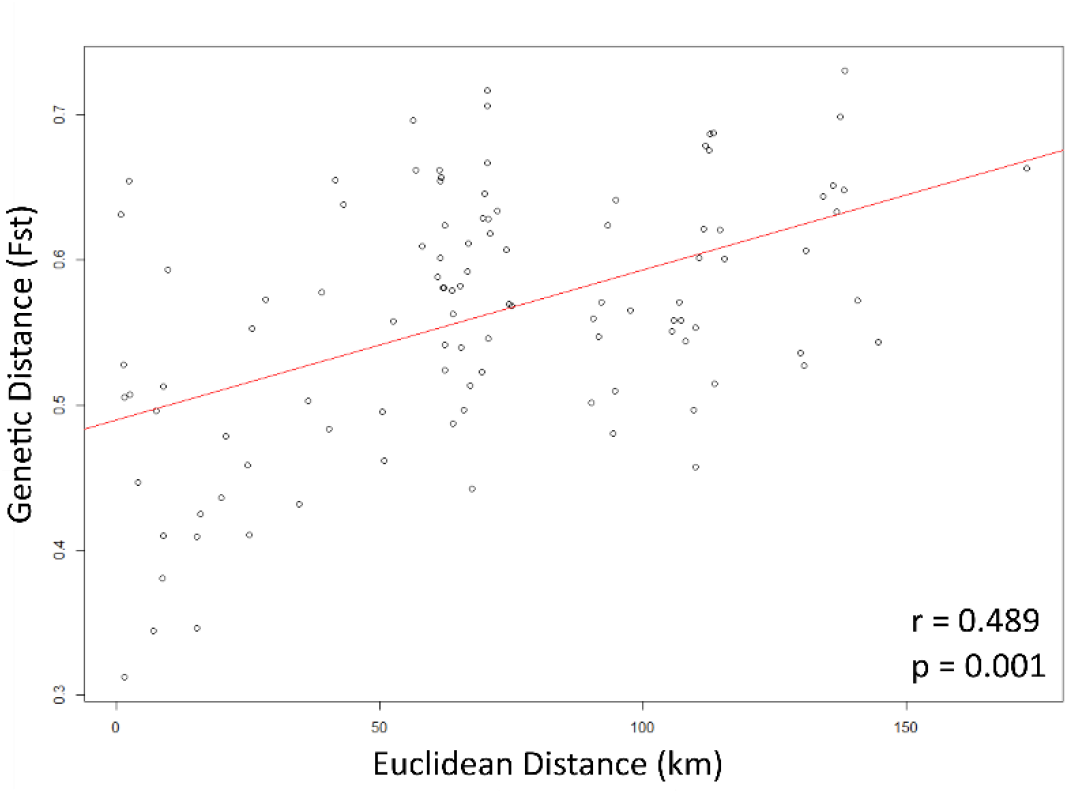
Scatterplot of genetic vs. geographic distance and outcome of statically significant partial Mantel test.

## Discussion

An understanding of diversification mechanisms in lichens has been hampered by a lack of population genetic studies in these diverse clades of fungi. This study is the first to report the results of a genomic approach for investigating the population structure of a lichen. Low-coverage whole-genome sequencing of lichen fragments produced large quantities of SNP data (>122,000 SNPs) among individuals within a species, even after contaminants were removed by stringent filtering. These results demonstrate that culturing is not required for lichen population genomics. The original hypothesis that the main dispersal strategy of *Cetradonia linearis* is through clonal propagation was not supported, as no clones were identified across or within sites. However, there is evidence that the species less frequently undergoes outcrossing and sexual recombination based on the high rates of linkage disequilibrium (~122K SNPs reduced to ~10K, Fig. 3), estimated selfing rates >0.5, and the putatively unisexual mating system.

In contrast to prior work suggesting fungi are not dispersal-limited, the hypothesis that there are low rates of gene flow among populations was supported by high F_st_ values (0.312–0.730), significant correlation between genetic and geographic distance (Mantel Test, r = 0.489, p < 0.01), and proportionally higher influence of geographic distance on genetic distance when compared to environmental distance (αE:αD < 1). There was no evidence for isolation by environment based on the partial Mantel test and BEDASSLE results. However, further studies of other environmental variables, such as average high temperatures of warm months or quarters, may reveal signals of adaptation not recovered here. Additionally, future spore trapping and viability assays would be a useful way to directly measure dispersal potential. The sites with the highest genetic diversity were concentrated in the southern portion of the range of the taxon as predicted, suggesting that these may have acted as refugial areas during the LGM. The results of this study support the notion that gene flow among fungal populations decreases with distance - populations show strong signs of IBD despite the production of small propagules (O’Malley 2007) - suggesting that genetic drift may represent a more important process in diversification of lichenized fungi than previously appreciated. Our data also indicate that recombination can be low despite the frequent presence of sexual spore producing structures. This finding highlights the phenomenon that observed reproductive mode does not necessarily translate directly to the frequency of recombination (Taylor et al. 2015).

### Influence of Reproductive Strategy on Population Genetic Structure in Lichens

Our data on *Cetradonia linearis* contribute to a growing understanding of the relationship between fungal reproductive types and genetic diversity and structure. Three species of lichenized fungi were previously examined with detailed population genetic studies including analysis of the mating-system (Itten and Honegger 2010; Singh et al. 2012; Alors et al. 2017). One of these species was investigated with RAPD-PCR and the other two with microsatellites, so comparisons among the studies, and with our study based on genomic data, must be done with consideration of the very different underlying data. Nonetheless, because there are no genomics studies on lichen population genetic structure currently published, a careful comparison of our results with this previous research is useful. The lichen *Xanthoria parietina* was found to be unisexual, having only the MAT1-2 gene present in all individuals investigated, and no observed instances of trichogynes, though it is almost always fertile (Scherrer et al. 2005). The population genetic structure of *X. parietina* based on RAPD-PCR fingerprinting revealed high rates of genotypic diversity within populations, even on a microsites scale, and much lower genetic diversity between populations than within them (Itten and Honegger 2010). A study of the lichen *Parmelina carporrhizans* found a similar pattern of very high gene flow among most populations sampled, though it is a heterothallic species (Alors et al. 2017). The pattern observed in these two species starkly contrasts with that of *Lobaria pulmonaria*, a heterothallic species that is often observed without sexual reproductive structures, in which apothecia usually are not produced until individuals are 15-25 years old (Denison 2003; Høistad and Gjerde 2011; Singh et al. 2012). *Lobaria pulmonaria* consistently displays high rates of clonality within populations and sampling sites (Werth et al. 2006; Sing et al. 2012). One way to explain the difference between the population genetic patterns of the two heterothallic species is the ratio of the two alternate MAT idiomorphs: *L. pulmonaria* ratios are often skewed in populations while *P. carporrhizans* populations have equal ratios (Singh et al. 2012; Alors et al. 2017). Population genetic structure and biology of *Cetradonia linearis* is more similar to *X. parietina* and *P. carporrhizans* because 1) it is almost always fertile, 2) no clones have been identified (defined as >95% shared SNPs for the genomic data), even from closely collected colonies, and 3) there is a high level of polymorphism within each population (π = 0.148 – 0.338). However, our results indicate *C. linearis* populations have low rates of gene flow, which contrasts with the pattern of little genetic structure found in both *X. parietina* and *P. carporrhizans.* Additional studies using genomics to investigate population genetics in lichenized fungi will allow more direct comparisons of genetic structure across diverse fungal clades. These results, along with the high rate of linkage disequilibrium, suggest that while *C. linearis* does not seem to frequently reproduce clonally, there must be some rate of self-fertilization or clonality and dispersal restriction that leads to the genetic isolation of populations. To draw large-scale conclusions about the influence of observed reproductive mode on recombination, further studies tackling a greater breadth of taxonomic sampling throughout lichenized fungi will be required. Specific efforts are needed to target phylogenetic, morphological, ecological, and reproductive diversity to determine what factors most strongly shape recombination.

### Biogeographic History

The southern Appalachian Mountain Range is one of the oldest continuously exposed land masses on earth, and has served as a refugium at multiple points in geological history (Braun 1950). Thus, though it is a relatively small area, the long and complex geological history of the region has shaped similarly strong, complex population genetics patterns in endemic species across multiple domains of life (Manos and Meireles 2015). The population genetics of *C. linearis* are no exception. The southern portion of the current extent of *C. linearis* was likely a refugium during Pleistocene glaciation. The lines of evidence to support this hypothesis include the observation that genetic diversity is higher in southern populations and location of suitable areas predicted by the hindcast SDM (Fig. 5). Interestingly, the model also suggested an expansion of the range to lower elevation areas (Fig. 5). This finding is consistent with hypotheses that ranges of present-day high-elevation endemics expanded downslope during Pleistocene glaciation (Crespi et al. 2003; Bruhl 1997; Premoli et al. 2007; Desamore et al. 2010). While this downslope expansion might have been expected to connect populations and diminish the signal of IBD, the data generated for this study still show a strong geographic signal of increasing structure with distance. Population genetic studies of other high-elevation, southern

**Figure 5.**
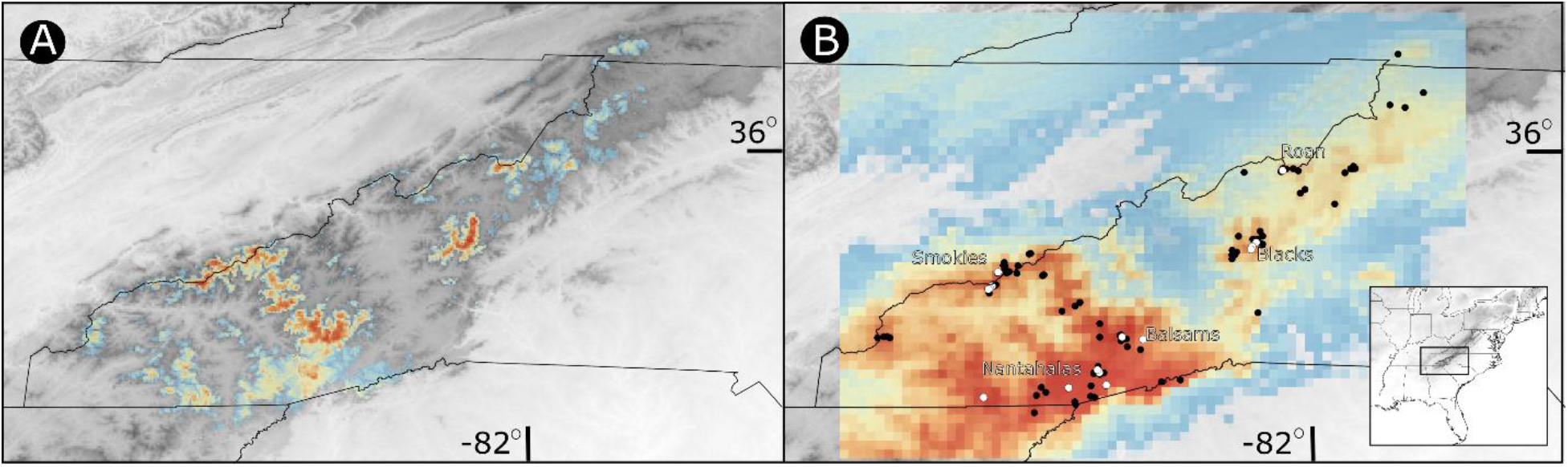
Species distribution model of *Cetradonia linearis* A) in the present; B) at the last glacial maximum. Probability of *C. linearis* grades from blue (low) to red (high). Inset gives larger spatial orientation of study area.

Appalachian endemics showed similarly strong signals of IBD, including the salamander *Desmognathus wrightii* (Crespi et al. 2003) and the spider *Hypochilus pococki* (Keith and Hedin 2012). A further parallel between the genetic structures of *D. wrightii* and *C. linearis* is that Roan Mountain populations did not group with populations from the Black Mountains, despite their close geographic proximity (Crespi et al. 2003). Population differentiation was so strong in *Hypochilus pococki* that the authors suggested it may actually be comprised of multiple cryptic species (Keith and Hedin 2012). The long and complex geological history of the southern Appalachians has resulted in not only high levels of species diversity, but also high genetic diversity within species.

## Conclusion

The results presented here provide strong evidence that the rare, narrowly endemic fungus *Cetradonia linearis* has highly geographically isolated populations over the small area of its distribution (Fig. 2). The population structure of *C. linearis* is congruent with other high-elevation southern Appalachian endemics, suggesting that dispersal among mountain peaks in the region is not frequent for multiple groups of organisms (Crespi et al. 2003; Keith and Hedin 2012; Fig. 5). We found no evidence of clones in our sampling, however we did find evidence for low rates of recombination, possibly facilitated by a homothallic reproductive system that allows self-fertility (Fig. 3). These results support a growing body of literature suggesting that fungal dispersal can be limited across relatively small spatial scales, despite the production of very small propagules (O’Malley 2007; Taylor et al. 2012). While other studies have found a strong influence for environmental factors influencing population structure of fungi (Branco et al. 2015), we found no evidence for isolation by environment (Supplement Table 1). Future comparative studies are required to full understand how extrinsic and intrinsic factors shape the population structure and recombination rates of fungi with different ecological requirements, life histories, and reproductive strategies (Grünwald et al. 2016; Peter and Schacherer 2016). These studies will be facilitated by rapid advancements in population genomics methods, which promise to reshape current perspective on fungal biology.

## Data Accessibility

(To be deposited upon publication)

Reference genome: Genbank Accession XXXX

Raw reads: Sequence Read Archive XXXX

## Acknowledgements

We would like to thank the following people for aiding with this research: Alex Cecil and Jenna Dorey for their field assistance, Gary Kauffman for help locating populations throughout the National Forest and many helpful discussions about *Gymnoderma*, Chris Ulrey for help locating the species on Blue Ridge National Park land, and especially for rapelling expertise, Dr. Richard Harris for conducting microscopy to search for trichogynes and spores. Library preparations and sequencing was conducted at The Rockefeller Genome Resource Center. Funding for this research came from Highlands Biological Station, NSF GRFP, and NSF DEB#1145511.

